# The *C. elegans* PTCHD homolog PTR-4 is required for proper organization of the pre-cuticular apical extracellular matrix

**DOI:** 10.1101/2021.03.30.437696

**Authors:** Jennifer D. Cohen, Carla E. Cadena del Castillo, Andres Kaech, Anne Spang, Meera Sundaram

**Author notes:** Corresponding Author: Meera Sundaram Department of Genetics University of Pennsylvania Perelman School of Medicine Philadelphia PA USA. Equal contributions (Jennifer D. Cohen, Carla E. Cadena del Castillo). Author contributions JDC, CECC, AS, and MS designed the experiments. CECC and JDC performed the experiments. AK generated the TEM data. CECC, JDC, AS and MS analyzed the data. JDC, AS, MS, and CECC wrote the manuscript, with input from all authors.

## Abstract

The Patched-related (PTR) superfamily of transmembrane proteins can transport lipids or other hydrophobic molecules across cell membranes. While the hedgehog receptor Patched has been intensively studied, much less is known about the biological roles of other PTR or Patched domain (PTCHD) family members. *C. elegans* has a large number of PTR/PTCHD proteins, despite lacking a canonical hedgehog pathway. Here, we show that PTR-4 promotes the assembly of the pre-cuticle apical extracellular matrix (aECM), a transient and molecularly distinct aECM that precedes and patterns the later collagenous cuticle or exoskeleton. *ptr-4* mutants share many phenotypes with pre-cuticle mutants, including defects in eggshell dissolution, tube shaping, alae (cuticle ridge) structure, and cuticle barrier function. PTR-4 localizes to the apical side of a subset of outward-facing epithelia, in a cyclical manner that peaks when pre-cuticle matrix is present. Finally, PTR-4 acts in a cell non-autonomous manner to properly localize the secreted ZP domain protein LET-653 to the pre-cuticle aECM. We propose that PTR-4 exports lipids or other hydrophobic components of the pre-cuticle aECM.

## Introduction

Patched (PTCH) and other related PTR/PTCHD proteins are members of a large, evolutionarily conserved family of small molecule and lipid transporters (Zhong et al., 2014). All are multi-pass transmembrane proteins with a predicted hydrophobic cleft to transport hydrophobic cargo or bind ligands, and all share structural homology with archeal and bacterial resistance-nodulation division (RND) transporters (Nikaido, 2018). Many organisms express multiple PTR proteins, which share a sterol sensing domain (SSD) and similar overall structure with classical PTCH proteins but show considerable sequence divergence (Kuwabara and Labouesse, 2002). While PTCH has been studied extensively for its role in canonical Hedgehog (Hh) signaling (Zhong et al., 2014), much less is known about the roles of other PTR proteins.

PTCH and several PTR proteins transport cholesterol across the plasma membrane or across other internal vesicle membranes. Although many details remain debated, recent evidence suggests that, in the absence of Hh signaling, PTCH transports cholesterol across the plasma membrane to locally inhibit the G-protein coupled receptor Smoothened (SMO). When Hh, a cholesterol-modified lipoprotein, binds the PTCH extracellular domain, it blocks cholesterol transport, allowing cholesterol to activate SMO signaling (Hu and Song, 2019; Petrov et al., 2020; Zhang et al., 2018). Another PTCHD protein, Dispatched, exports Hh from cells (Hall et al., 2019). Yet another related protein, Npc1, exports cholesterol from lysosomes; mutations in *Npc1* lead to massive, cholesterol-filled lysosomes and severe cellular dysfunction (Cologna and Rosenhouse-Dantsker, 2019; Pfeffer, 2019). In bacteria, some RND transporters export glycolipids and other hydrophobic cargo into the periplasm to aid in outer cell envelope biogenesis (Melly and Purdy, 2019). Other RND transporter family members pump metals or antibiotics (Nikaido, 2018). Despite the wide variation in these family members’ functions, they seem primarily to act as pumps for hydrophobic cargo.

In addition to PTCH, Dispatched, and Npc1, mammals express four different PTCHDs whose molecular functions are unknown, but which have intriguing possible links to disease. While they may not play a role in Hh signaling, at least PTCHD2 has been linked to maintenance of cholesterol homeostasis (Konířová et al., 2017; Zikova et al., 2009). Loss of and mutations in PTCHD1 are associated with autism spectrum disorders (Chaudhry et al., 2015; Torrico et al., 2015; Ung et al., 2018). PTCHD4 appears to negatively regulate Hh signalling in cancer (Chung et al., 2014), while the function of the non-essential PTCHD3 remains elusive.

The roundworm *C. elegans* contains two PTCH proteins (PTC-1,-3) and a large number of other PTR/PTCHD proteins, along with many unusual Hh-like proteins, but it lacks SMO and other members of the canonical Hh signaling pathway (Bürglin, 2008; Bürglin and Kuwabara, 2006; Hao et al., 2006b; Zugasti et al., 2005). We showed recently that PTC-3 has cholesterol transport activity (Cadena del Castillo et al., 2019). *C. elegans* may therefore be an excellent model for studying the broader transport roles for PTR/PTCHD proteins independent of SMO signaling.

Previous studies showed that RNAi-mediated knockdown of many *C. elegans ptr* genes led to defects in cuticular structure and molting (Hao et al., 2006a; Zugasti et al., 2005), suggesting a potential role for PTR proteins in trafficking or export of components of the apical extracellular matrix (aECM). *C. elegans* has multiple distinct types of aECM, including a chitinous eggshell and a collagenous cuticle that must be disassembled and rebuilt during each of four larval molts (Cohen and Sundaram, 2020; Page and Johnstone, 2007; Stein and Golden, 2018). Prior to the generation of each new cuticle, a molecularly distinct aECM, termed the sheath or pre-cuticle, coats external epithelia (Cohen et al., 2020b; Cohen and Sundaram, 2020; Priess and Hirsh, 1986; Vuong-Brender et al., 2017). Similar to some mammalian aECMs such as lung surfactant (Pérez-Gil, 2008) or eye tear film (Dartt, 2011), all of these *C. elegans* aECMs appear to contain layers with significant lipid content (Bai et al., 2020; Blaxter, 1993; Dartt, 2011; Forman-Rubinsky et al., 2017; Pu et al., 2017; Schultz and Gumienny, 2012). We were therefore intrigued by the possibility that PTR proteins might transport lipids or other hydrophobic cargos into one or more of these aECMs.

Here, we report that one *ptr* gene, *ptr-4,* is required for pre-cuticle aECM assembly. The pre-cuticle is a transient aECM that contains multiple proteins of the Zona Pellucida (ZP) domain, extracellular leucine-rich repeat only (eLRRon), and lipocalin (putative lipid transporter) families (Cohen et al., 2019; Cohen et al., 2020b; Forman-Rubinsky et al., 2017; Gill et al., 2016; Katz et al., 2018; Kelley et al., 2015; Mancuso et al., 2012; Vuong-Brender et al., 2017). It is present in the developing embryo and prior to each larval molt, and it shapes multiple epithelial tissues and helps pattern the subsequent collagenous cuticle (Cohen and Sundaram, 2020). We show that *ptr-4* mutants share many phenotypes with pre-cuticle mutants, including defects in eggshell dissolution, tube shaping, alae (cuticle ridge) structure, and cuticle barrier function. PTR-4 localizes to apical surfaces of outward-facing epithelia during periods of pre-cuticle assembly and remodeling, and PTR-4 acts in a cell non-autonomous manner to properly localize the secreted ZP domain protein LET-653 to the pre-cuticle aECM. Together, these data point to a role for PTR-4 in the transport of pre-cuticle aECM components.

## Results

### PTR-4 is essential for embryo hatching and for maintenance of lumen diameter within the excretory duct tube

To define *ptr-4* functions, we examined *ptr-4* mutants. Using CRISPR/Cas9 (Methods), we generated a new putative null allele of *ptr-4, cs273.* This allele has a 23 bp deletion that disrupts the first predicted transmembrane domain of PTR-4, followed by a frame shift and early stop (Fig 1A). We also obtained another deletion allele, *ok1576*, from the knockout consortium (Moerman and Barstead, 2008); *ok1576* contains a 2.2 kb in-frame deletion that removes most of the Patched domain but leaves the first and last transmembrane domains intact (Fig 1A). Both alleles were 100% embryonic lethal (Fig 1B). Both were efficiently rescued by a large genomic fragment containing the *ptr-4* gene (Fig 1B), indicating that lethality was indeed caused by loss of *ptr-4*.

**Figure 1.**
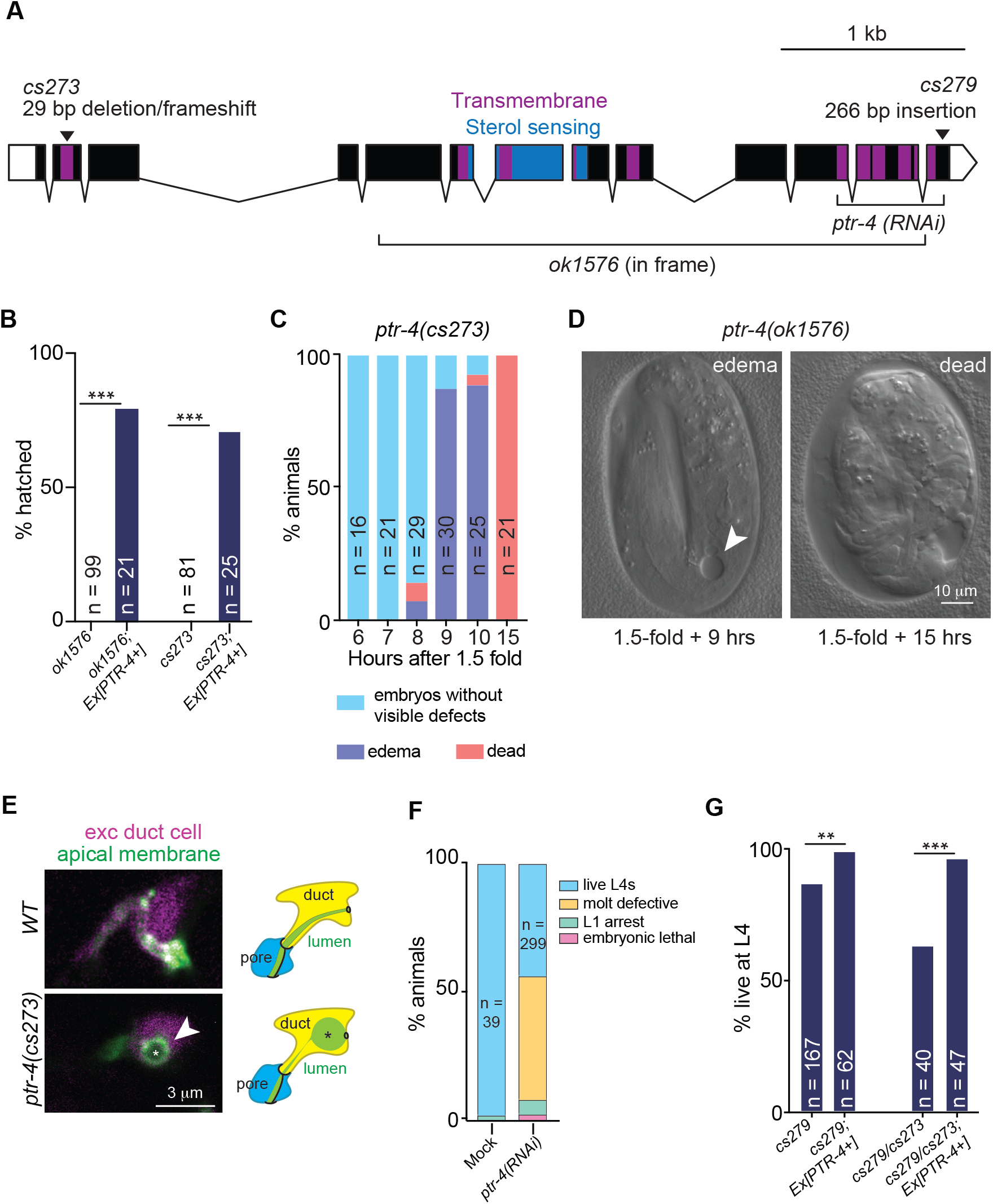
*ptr-4* null mutants are embryonic lethal. (A) *ptr-4* gene model with the transmembrane domains indicated in purple. Five of the twelve predicted TM domains comprise the sterol sensing domain (blue), a domain also found in some non-PTR proteins that bind cholesterol (Kuwabara and Labouesse, 2002). The studied mutations and target of the RNAi are indicated. *ok1576* is a 3093 bp in-frame deletion that removes the sterol sensing domain and 9 transmembrane domains. We used CRISPR/Cas9 to generate *cs273*, which is a 29 base pair deletion causing a frameshift and early stop within the first transmembrane domain (see also Figure S1). *cs279* is a 266bp insertion at the *ptr-4* 3′ end containing unrelated sequences derived from the genes *rol-6* and *pie-1* (see also Figure S1). *cs279* is predicted to delete the last three amino acids of PTR-4 and replace them with seventeen unrelated amino acids before a new stop codon that is then followed by 213 additional nucleotides before the endogenous stop codon; this extended 3′ UTR may interfere with normal gene expression. (B) *ptr-4* mutants of either allele failed to hatch and were rescued by a large genomic fragment containing the normal *ptr-4* gene. *** P < 0.0001, Fisher’s exact test. (C) In *ptr-4(cs273)* mutants, edemas form in the hours after the 1.5-fold stage, and eventually the embryos die without hatching. (D) Differential interference contrast (DIC) images of *ptr-4(ok1596)* embryos at the indicated timepoints. Arrowhead indicates edema. (E) Edema formed within the excretory duct tube lumen. Confocal images of *wild type* (*WT*) and *ptr-4(cs273)* mutants with the duct cytoplasm marked with *lin-48pro::mRFP* and the apical membrane marked with *RDY-2::GFP*. Schematic interpretations of the phenotype are shown at right. (F) *ptr-4(cs279)* is a hypomorph. Many *cs279* mutants survive but larval lethality is enhanced in *cs279/cs273* trans-heterozygotes. To assess trans-heterozygote survival, *cs279* males were crossed to *cs273*; *Ex[PTR-4+]* hermaphrodites and their non-transgenic progeny were scored for lethality. *cs279* lethality is rescued by a large genomic fragment containing *ptr-4*. *** P < 0.0001, ** P < 0.001, Fisher’s exact test. (G) Many *ptr-4(RNAi)* larvae died due to incomplete molting with small fractions dying as very small L1 larvae (L1 arrest) or as embryos.

Upon closer inspection, we found that *ptr-4* mutant embryos had defects in hatching and in maintaining lumen diameter within tubes of the excretory system. *ptr-4* mutant embryos appeared normal until the time period when they should hatch (Fig. 1C). Hatching never occurred, however, suggesting a defect in eggshell dissolution. Embryos subsequently developed edemas that began as large dilations within the lumen of the excretory duct tube, a narrow unicellular tube that is required for fluid excretion (Fig. 1C-E). By 15 hours after the expected time of hatch, all embryos were dead (Fig. 1C,D). Notably, both the hatching defect and excretory phenotypes of *ptr-4* mutants resemble defects observed in mutants lacking the lipocalin LPR-3 (a putative lipid transporter) or various other known pre-cuticle aECM components that line the epidermis and/or the duct tube channel (Cohen et al., 2019; Forman-Rubinsky et al., 2017; Gill et al., 2016; Hishida et al., 1996; Mancuso et al., 2012; Pu et al., 2017; Stone et al., 2009).

### PTR-4 is required for proper cuticle patterning and permeability barrier formation

To examine requirements for *ptr-4* at stages later than embryogenesis, we turned to RNAi and to a hypomorphic allele, *cs279. ptr-4(RNAi)* caused relatively minor embryonic or early larval lethality, but about 50% of the animals had molting defects similar to those described previously (Zugasti et al., 2005) (Fig. 1F). The remaining animals progressed to adulthood but were small (Fig. 2A), uncoordinated (Fig. 2B), and showed cuticle defects as described below. In *C. elegans*, locomotion requires alternating contraction of body wall muscles that are connected to the cuticle (Page and Johnstone, 2007). In addition, the cuticle has to maintain osmolarity and turgor pressure to consequently achieve movement (Moribe et al., 2004). It is likely therefore that cuticle defects contribute to the observed defects in locomotion.

**Figure 2.**
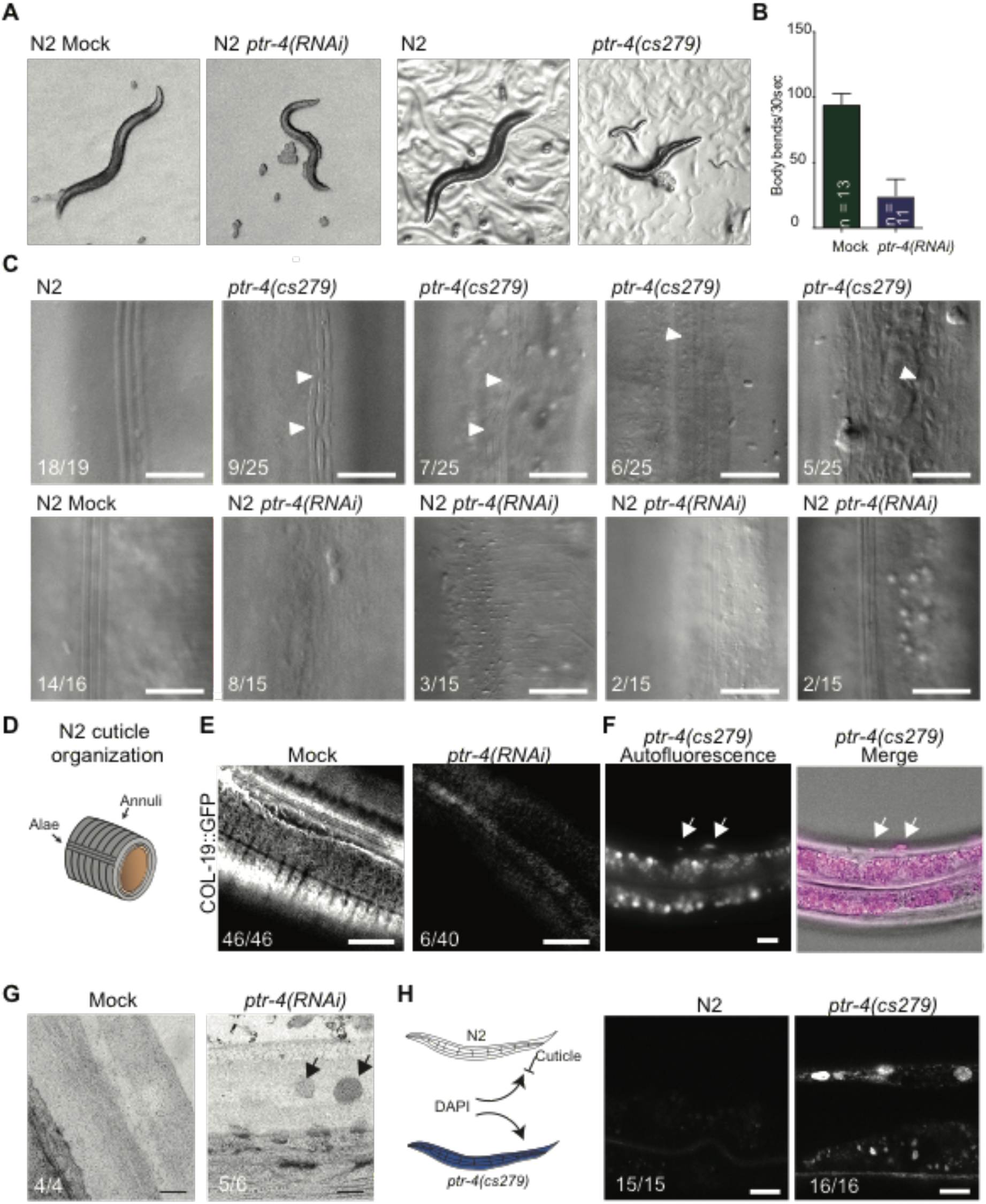
PTR-4 depletion induces cuticular defects. (A) *ptr-4(cs279)* and *ptr-4(RNAi)* adults are small and uncoordinated. (B) Mock treated and *ptr-4(RNAi)* worms were placed in M9 media and were allowed to swim. Body bend analysis in a 30 seconds test showed reduced motility upon *ptr-4(RNAi)*. (C) DIC images of N2 and *ptr-4(cs279)* worms. The cuticle in *ptr-4(cs279)* and *ptr-4(RNAi)* worms is structurally impaired with multiple types of alae defects. (D) Schematic depiction of the cuticle with annuli and alae. (E) PTR-4 loss disrupts COL-19::GFP localization. COL-19::GFP marked annuli in all wild-type adults, but was detectable in only 6/40 *ptr-4(RNAi)* adults. (F) *ptr-4(cs279)* animals have abnormal packets of autofluorescent material within the cuticle. (G) TEM images showing cuticle structure in Mock and *ptr-4(RNAi)* treated animals. In *ptr-4(RNAi)* animals, some vesicular-like structures were observed within the cuticle. Scale bar, 150 nm. (H) Permeability test using the impermeable dye DAPI. In animals with an intact cuticle DAPI, is not able to infiltrate and stain cell nuclei. *ptr-4(RNAi)* makes the cuticle permeable. Representative DAPI images of N2 or *ptr-4(cs279)* animals upon incubation with DAPI. Scale bars 10 μm.

We inadvertently generated the allele *ptr-4(cs279)* when attempting to tag the locus using CRISPR/Cas9 (Methods). *cs279* contains a 266 bp insertion of DNA from the genes *rol-6* and *pie-1* inserted just before the *ptr-4* stop codon, leading to modification of the protein and extension of the 3’ UTR (Fig. 1A and Fig. S1). Most *ptr-4(cs279)* animals survived (Fig. 1G) but were small and uncoordinated, resembling the *ptr-4(RNAi)* adults above (Fig. 2A). To confirm that *cs279* is hypomorphic, we demonstrated that its mild lethality is rescued by a large genomic fragment containing *ptr-4* (Fig.1G). Next, we generated compound heterozygotes that contained the *ptr-4(cs279)* allele over the *ptr-4(cs273)* null allele. These worms had increased post-embryonic lethality compared to *cs279* homozygotes (Fig. 1G). Together, these data indicate that *cs279* is a mild hypomorphic allele.

Surviving *ptr-4(cs279)* and *ptr-4(RNAi)* adults had abnormal cuticle structure. The adult epidermal cuticle is organized into circumferential ridges (annuli) that cover the bulk of the epidermis, and longitudinal ridges (alae) that cover the lateral seam cells (Page and Johnstone, 2007) (Fig. 2C,D). Animals with reduced *ptr-4* activity displayed a variety of alae defects, ranging from interrupted alae to almost complete absence of alae (Fig. 2C). Furthermore, the adult-specific collagen COL-19::GFP (Thein et al., 2003) was depleted or lost from the annuli (Fig. 2E). By light microscopy, packets of autofluorescent material were observed within the epidermal cuticle (Fig. 2F). These abnormal structures in *ptr-4(RNAi)* animals were also observed by transmission electron microscopy (Fig. 2G), and closely resembled structures previously reported for mutants of the pre-cuticle factor LET-4 (Mancuso et al., 2012). Finally, we found that, similar to many pre-cuticle or cuticle mutants (Cohen et al., 2020b; Forman-Rubinsky et al., 2017; Mancuso et al., 2012; Sandhu et al., 2021), the cuticle in *ptr-4(RNAi)* animals was permeable to small molecules (Fig. 2H), indicating that the permeability barrier was defective.

### PTR-4 localizes transiently to the apical plasma membrane of outward-facing epithelia

To determine when and where PTR-4 localizes, we used CRISPR/Cas9 to tag endogenous PTR-4 with the fluorescent protein Superfolder (Sf) GFP (Pedelacq et al., 2006) (Fig. 3A). PTR-4::SfGFP was not detected during the early part of embryogenesis but appeared by the 2-fold stage along the apical membranes of the epidermis, excretory duct and pore, and rectum (Fig. 3B,C). However, PTR-4 was transient and rapidly cleared prior to hatching (Fig. 3B,D). PTR-4 reappeared in the middle of each subsequent larval stage, during the time when pre-cuticle is present (Cohen and Sundaram, 2020; Forman-Rubinsky et al., 2017; Gill et al., 2016). During L4 stage, the stage preceding adulthood, PTR-4 was present on apical surfaces between the major epidermis (hyp7) and the lateral seam cells (Fig. 3E), which secrete alae (as well as pre-cuticle factors that are needed to shape alae (Cohen et al., 2019; Flatt et al., 2019; Forman-Rubinsky et al., 2017; Kolotuev et al., 2009; Liégeois et al., 2006)). PTR-4 was also present along apical surfaces of the rectum and of some cells in the vulva, the tube through which eggs will be laid (Fig. 3E). As in embryogenesis, PTR-4 was transient and disappeared as the adult cuticle was made. Together, these observations demonstrate that PTR-4 is present on the apical surfaces of external epithelia during the time period when pre-cuticle is present.

**Figure 3.**
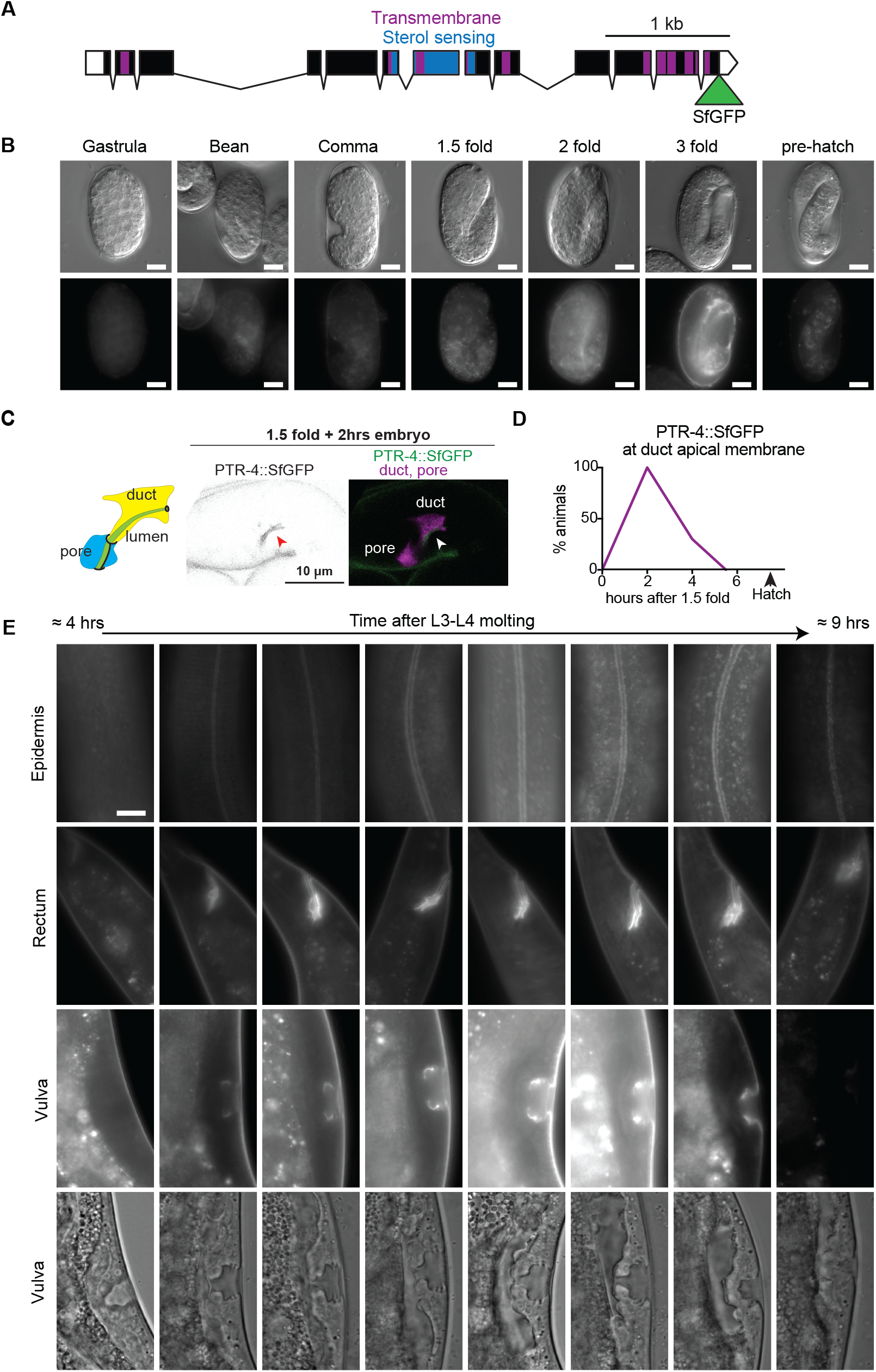
PTR-4 localizes transiently to apical surfaces of external epithelia. (**A**) *ptr-4* gene model with the transmembrane domains indicated in purple and sterol sensing domain in blue. Superfolder GFP (SfGFP) was inserted into the *ptr-4* endogenous locus via CRISPR/Cas9 technology to make *ptr-4(cs264 [PTR-4::SfGFP])*. (B) PTR-4::SfGFP localization during worm embryonic development. PTR-4::SfGFP first becomes visible at the 2-fold stage in the epidermis, rectum, and excretory duct/pore and then disappears before hatching. Scale bar 10 μm. (C) PTR-4::SfGFP localizes to the apical surface of the excretory duct tube (arrowhead) at 1.5-fold + 2hrs embryonic stage. For visualization, the duct and pore cells were labeled with *grl-2pro::mRFP*. (D) Time course of PTR-4::SfGFP expression on the duct apical membrane (n = 20 per timepoint). (E) PTR-4::SfGFP localization in L4 sub-staged larva (n=5 or more per timepoint). PTR-4::SfGFP expression in the epidermis, rectum, and vulva was dynamic during development. While in early L4, there is no PTR-4::SfGFP signal, in mid L4 it can be observed at the plasma membrane. In late L4 animals, PTR-4::SfGFP localizes to internal structures and prior to molting the signal disappears. Scale bars 10 μm.

### PTR-4 transient plasma membrane localization is regulated by oscillating protein translation and endocytosis

Cellular protein concentration is dependent on protein synthesis and degradation. To investigate the mechanisms controlling PTR-4 plasma membrane expression, we first analyzed published transcriptional profiling and ribosomal footprint data during larval development (Hendriks et al., 2014). Both *ptr-4* mRNA levels and ribosome occupancy oscillated during development with peaks during intermolts and troughs during molting (Fig. 4A). This oscillatory behavior is also shared by most aECM-encoding genes, with pre-cuticle factors (e.g. LET-653) and some of the earliest cuticle collagens peaking slightly before or at the same time as PTR-4, and most cuticle collagens peaking later (Fig. 4A).

**Figure 4.**
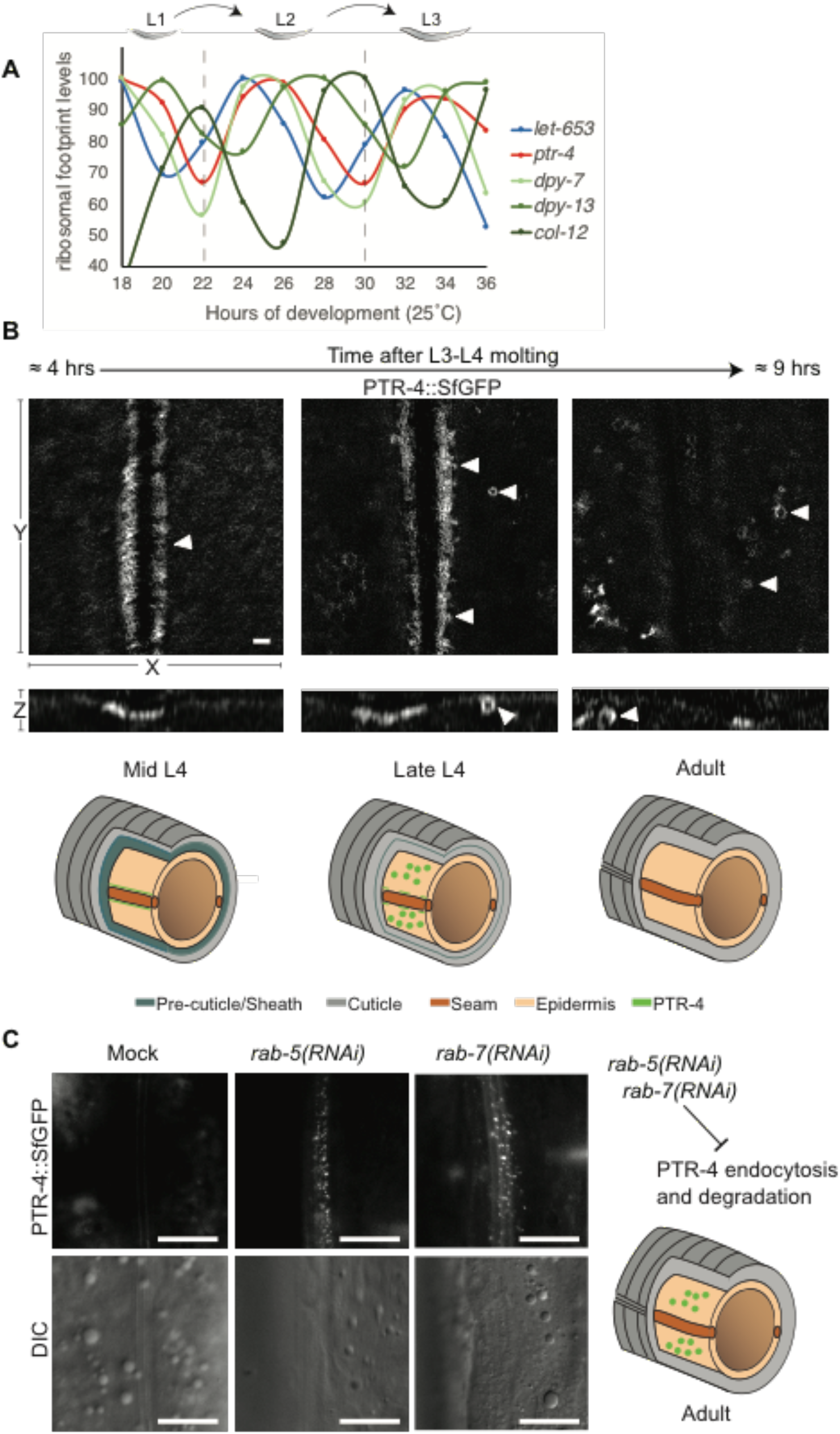
PTR-4 dynamic localization is regulated by endocytosis. (A) PTR-4 ribosomal footprints oscillate during larval development, with peaks coinciding with those of the earliest cuticle collagens. LET-653, DPY-7, DPY-13 and COL-12 are shown as representatives of pre-cuticle and early, intermediate, or late collagens, respectively (Page and Johnstone, 2007). Data are extracted from a previously published dataset (Hendriks et al., 2014). (B) Structural illumination (SIM) images of PTR-4::SfGFP in the epidermis. In mid-L4 worms, PTR-4::SfGFP is present at the hyp7 plasma membrane (n=6). As development proceeds, PTR-4::SfGFP gathers in membrane invaginations (n=6). At late L4 stages, PTR-4::SfGFP localizes exclusively in intracellular compartments (n=5). Scale bar 1 μm. Schematics below (adapted from Katz et al 2018) summarize the observed changes in PTR-4 localization. (C) PTR-4 localization changes during development are due to endocytosis. RNAi of *rab-5* (n=17/19) or *rab-7* (n=13/14) blocked endocytosis and/or degradation, leading to accumulation of PTR-4 within the epidermis of adult worms, whereas 0/10 mock-treated animals had detectable PTR-4 at this stage.

Next, we determined how PTR-4 degradation might be regulated. To this end, we aimed to follow changes in PTR-4 plasma membrane localization in the epidermis, using structured illumination microscopy (SIM) (Fig. 4B). After the L3-L4 molt, PTR-4 was localized at the apical plasma membrane of hyp7, facing the seam cells. In mid-L4, PTR-4 started to move away from the contact site with the seam cells and sites of PTR-4 endocytosis were observed. At the early adult stage, PTR-4 was absent from the contacts between epidermis and seam cells. If PTR-4 was indeed endocytosed and then degraded in the lysosome, interfering with this pathway should stabilize PTR-4. When we knocked down the key endocytic regulators RAB-5 and RAB-7, PTR-4 still moved away from the epidermis-seam cell contact but was no longer efficiently endocytosed and degraded (Fig. 4C).

In summary, our data indicate that PTR-4 transient plasma membrane expression is regulated through oscillating PTR-4 protein synthesis on one hand and endocytosis and lysosomal degradation on the other hand.

### PTR-4 functions cell non-autonomously to properly localize the ZP protein LET-653 in the vulva pre-cuticle

The similarity of *ptr-4* mutants to known pre-cuticle mutants and the spatial and temporal pattern of PTR-4 localization are consistent with a possible role in pre-cuticle assembly or remodeling. Furthermore, the SIM data above also showed that PTR-4 is expressed on epidermal cells adjacent to seam cells, rather than on the alae-producing seam cells themselves, suggesting that PTR-4 can act cell non-autonomously to affect aECM assembly.

To further test this hypothesis, we turned to the vulva tube, where different combinations of pre-cuticle factors assemble on the apical surfaces of each different cell type (Cohen et al., 2020b). We showed previously that, during mid-L4, the secreted zona pellucida (ZP) domain of LET-653 specifically decorates the apical surfaces of the dorsal-most vulE and vulF cells, which appear to have unique properties conducive to LET-653(ZP) assembly (Cohen et al., 2020a; Cohen et al., 2020b) (Fig. 5A,B). However, when PTR-4 was knocked-down by RNAi, LET-653(ZP)::SfGFP was no longer concentrated at these surfaces, but instead filled the entire vulva lumen (Fig. 5A,B). There appeared to be no defect in LET-653 secretion; however, the assembly of LET-653(ZP) in the membrane-proximal ECM was defective. Importantly, PTR-4::SfGFP was never observed on vulE or vulF cells, but rather marked the apical membranes of the more ventral cells (Fig. 5C). Vulva cell type specificity was further confirmed by examining *lin-12/Notch* mutants with cell fate alternations (Fig. 5C). Based on the inversely correlated expression patterns of PTR-4 and LET-653(ZP), we infer that PTR-4 does not directly anchor LET-653(ZP) or other local aECM factors, but must act cell non-autonomously to promote their proper assembly (Fig. 5D).

**Figure 5:**
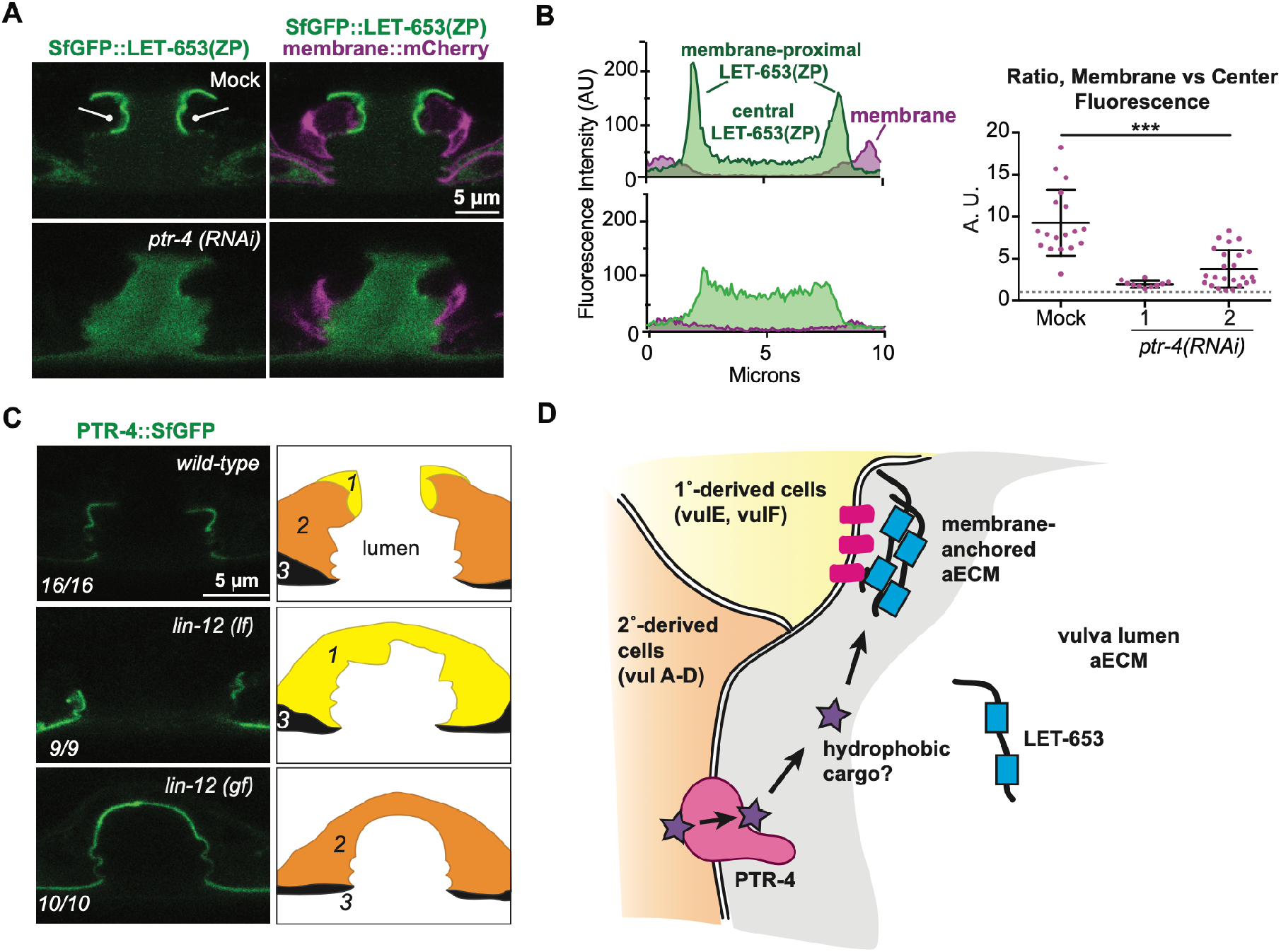
PTR-4 is required for proper assembly of the vulva pre-cuticle matrix. (A) Localization of LET-653(ZP) to vulE and vulF cell surfaces is dependent on PTR-4. Knockdown of PTR-4 does not impair secretion of LET-653(ZP), but the integration into the ECM is impaired. (B) Quantification of data shown in (A). Control n=18; *ptr-4(RNAi)* 1 n=10. *ptr-4(RNAi)* 2 n = 24. *** P < 0.0001, Mann-Whitney U-test. (C) PTR-4::SfGFP is expressed at apical membranes of cells derived from 2° or 3° lineages, and not in vulE/F cells derived from 1° lineages. *lin-12/Notch* mutations were used to alter cell fates as indicated in schematics (Greenwald et al., 1983). (D) Model for non-autonomous effect of PTR-4 on aECM assembly. PTR-4 may transport a hydrophobic cargo that non-cell-autonomously aids in aECM assembly of LET-653.

## Discussion

The existence of so many *C. elegans* PTR family members has been a long-standing puzzle, given the absence of a canonical Hedgehog signaling pathway. Recent evidence for the regulation of plasma membrane cholesterol levels by PTCH, combined with the homology to bacterial RND transporters, suggests a unifying hypothesis that all PTCHD/PTR proteins are transmembrane pumps for hydrophobic cargos. However, the biological roles that these proteins fulfill remain little characterized. Prior work suggested that many *C. elegans* PTR proteins affect development of the collagenous cuticle (Hao et al., 2006a; Zugasti et al., 2005). Here we provided evidence that *C. elegans* PTR-4 promotes proper assembly of the pre-cuticle, a molecularly distinct aECM that precedes and patterns the cuticle. First, *ptr-4* mutants display a set of phenotypes that closely resemble those of known pre-cuticle mutants. Second, the presence of PTR-4 on apical membranes suggests a role in transporting a small hydrophobic molecule across those membranes, and its oscillatory temporal pattern and restriction to external epithelia is consistent with a role in transporting factors relevant to pre-cuticle synthesis. Third, PTR-4 is required for the assembly (though not secretion) of at least one specific pre-cuticle aECM factor, the ZP protein LET-653. Finally, we note that there is prior evidence for a role of the PTR DAF-6 in organizing the aECM of glial sheath and socket tubes (Perens and Shaham, 2005). Given the varied types of aECM that are present on different epithelial surfaces and at different stages of the *C. elegans* life cycle (Cohen and Sundaram, 2020), it is plausible that the large family of *C. elegans* PTR proteins are dedicated to transporting different types of lipids and small molecules that contribute (directly or indirectly) to these diverse aECMs.

### Relationship of PTR-4 to the eggshell, pre-cuticle, and cuticle aECMs

The pre-cuticle is a transient aECM that covers developing epithelia prior to and during early stages of cuticle collagen deposition and maturation (Cohen and Sundaram, 2020). The pre-cuticle also sits at the interface of the embryo and the inner eggshell. The *ptr-4* mutant phenotypes described here suggest defects in all three of these distinct aECMs (the eggshell, pre-cuticle, and cuticle). This could reflect multiple distinct roles for PTR-4, or all of the defects could be explained by a common role, such as a role in pre-cuticle assembly and remodeling.

*ptr-4* and *lpr-3* are not expressed until long after the eggshell forms, and mutants do not exhibit early developmental defects characteristic of eggshell mutants (Stein and Golden, 2018), so failure to dis-assemble the eggshell in these mutants is unlikely to reflect a problem in the main eggshell structure itself. Potentially, the innermost layer of the eggshell could be impacted by the aberrant formation of the pre-cuticle matrix beneath it, leading to hatching defects. Alternatively, eggshell dis-assembly may require specific “I’m ready” signals to be released from the embryonic cuticle, and aberrant cuticle organization could disrupt this communication. Finally, *ptr-4* could be required to synthesize or transport a relevant signal, such as a steroid hormone that triggers hatching. Normal hatching in all known cuticle mutants and the clearance of PTR-4 prior to hatching (Fig. 3) make the second two models less likely. The mechanisms that control hatching in *C. elegans* are largely unknown, and *ptr-4* and *lpr-3* should be useful tools for further studies.

We suggest that *ptr-4* cuticle defects could also be an indirect consequence of aberrant pre-cuticle, since the pre-cuticle patterns the cuticle (Cohen and Sundaram, 2020). Defects in LET-653(ZP) localization clearly demonstrate that *ptr-4* loss disrupts the pre-cuticle. Loss of LET-653 alone would not be sufficient to explain all of the observed phenotypes, but many other pre-cuticle mutants exhibit *ptr-4*-like defects in excretory duct tube collapse/dilation, alae structure, permeability, or molting. In contrast, such defects are seen more rarely or not at all in cuticle collagen mutants, most of which have relatively mild phenotypes. Notably, a recent survey of cuticle collagens found that only six of the earliest expressed collagens (DPY-2, DPY-3, DPY-7, DPY-8, DPY-9, and DPY-10) are required for maintaining cuticle barrier function (Sandhu et al., 2021). A subset of these have also been implicated in alae formation (McMahon et al., 2003). These “pioneer collagens” are expressed during the time that pre-cuticle (and PTR-4) are present (Fig. 4A) (Hendriks et al., 2014), so defects in pre-cuticle could secondarily affect their deposition or patterning. The process by which the pre-cuticle is exchanged for the cuticle is another area that is poorly understood and needs further study.

### Models for PTR-4 function in pre-cuticle organization

Depending on their topology, PTR proteins can transport cargo away from or towards the cell cytoplasm. For example, Dispatched exports Hh out of cells and into the environment (Hall et al., 2019), while NPC exports cholesterol out of the lysosome lumen and into the cytoplasm (Cologna and Rosenhouse-Dantsker, 2019; Pfeffer, 2019). The directionality with which Patched transports cholesterol is still debated in the literature (Hu and Song, 2019; Petrov et al., 2020; Zhang et al., 2018), though most data, including our data with *C. elegans* PTC-3 (Cadena del Castillo et al., 2019), suggest a cholesterol export function. We unfortunately failed to achieve good expression of PTR-4 in either *S. cerevisiae* or HEK293 cells, making it impossible to check directly for extrusion of cholesterol or other lipid types by PTR-4; therefore, its specific cargo and direction of transport remain unknown. Given the unusually large number of PTR proteins in *C. elegans*, along with the diverse cargos known for related bacterial RND transporters (Nikaido, 2018), it is likely that different PTR proteins transport different classes of cargo.

Given the above, combined with the primary location of PTR-4 at apical plasma membranes, the connection of PTR-4 with the pre-cuticle aECM could be at least threefold. First, PTR-4 could regulate the plasma membrane lipid composition in its vicinity, thereby either promoting or reducing the resident time of aECM remodelers at the plasma membrane and/or negatively or positively regulating the secretion of ECM components. Second, PTR-4 could import small lipophilic molecules into cells, possibly helping to clear aECM components or to provide building blocks for aECM production or trafficking. Third, PTR-4 could extrude lipids or other small lipophilic molecules that are either regulators or actual components of the aECM. For example, candidate PTR-4 cargoes include members of the divergent Hh-like family in *C. elegans,* many of which also are required for proper aECM organization (Bürglin, 2008). The apparently cell non-autonomous effects of PTR-4 on LET-653(ZP) assembly in the vulva would favor a model whereby PTR-4 mediates export of factors from 2° and 3° lineage-derived cells that ultimately impact LET-653 assembly on the 1° lineage-derived vulE/F cells (Fig. 5D).

A role for PTR-4 in aECM export would be analogous to the roles of MmpL (mycobacterial membrane protein large) proteins, which export various complex glycolipids across the inner cell membrane and into the periplasmic space for eventual incorporation into the outer cell membrane or other layers of the mycobacterial cell envelope (Melly and Purdy, 2019). Although the lipid content of the pre-cuticle aECM is not known, many aECMs, including the *C. elegans* eggshell and cuticle, do have significant lipid content, which is thought to confer their permeability barrier functions (Bai et al., 2020; Blaxter, 1993; Cohen et al., 2020b; Olson et al., 2012; Schultz and Gumienny, 2012). When viewed by TEM, the pre-cuticle aECM has an outer, electron-dense layer that could serve as a lipid-rich covering to corral developing membrane-proximal aECM layers and keep them separate from the more dynamic matrix in the central part of tube lumens (Cohen et al., 2020b; Gill et al., 2016; Mancuso et al., 2012). Loss of such a layer could allow secreted factors like LET-653 to move away from the membrane too quickly, hampering their local assembly (Fig. 5D).

Another clue to the importance of lipids in the pre-cuticle aECM is the requirement for multiple lipocalins, which are secreted transporters of small lipophilic molecules. The lipocalin LPR-3 directly incorporates into the pre-cuticle aECM, and *lpr-3* mutants share hatching, excretory tube, permeability, and molting phenotypes with *ptr-4* mutants (Forman-Rubinsky et al., 2017). Another lipocalin, LPR-1, does not incorporate into the pre-cuticle, but acts cell non-autonomously to affect its organization or function (Forman-Rubinsky et al., 2017; Pu et al., 2017). Potentially, PTR-4 and lipocalins could act in tandem to transport lipophilic cargoes across membranes and through extracellular spaces. Further tests of this idea will require identification of the relevant cargos.

## Materials and Methods

### Worm strains, alleles, and transgenes

All strains were derived from Bristol N2 and were grown at 20° under standard conditions (Brenner, 1974). *ptr-4(cs273)* mutant embryos were obtained from mothers rescued with a *ptr-4*(+) transgene, *csEx888*. This transgene was generated by coinjecting fosmid WRM069dE04 at 15 ng/μl along with pIM175 (unc-119::gfp) at 30 ng/μl and SK+ at 100 ng/μl. See Table S1 for a complete list of strains and alleles used.

### *ptr-4* mutant isolation and allele identification

The *ptr-4(cs264[PTR-4::SfGFP]), ptr-4(cs273)* and *(cs279)* alleles were created in an N2 background by CRISPR-Cas9. *ptr-4(cs264[PTR-4::SfGFP])* was generated using the SapTrap method (Schwartz and Jorgensen, 2016). Briefly, *ptr-4* single-guide RNA (TTTCTAGAGTCGGGGTGAAA) was inserted into the Cas9 expression plasmid pDD162 (Dickinson et al., 2013) and co-injected with a plasmid (pJC59) containing *ptr-4* homology arms attached to SfGFP derived from the plasmid pMLS279 (Schwartz and Jorgensen, 2016). *ptr-4(cs264[PTR-4::SfGFP])* animals were viable and appeared normal (n=93/96 normal, 3/96 unaccounted for), indicating the tag does not interfere with function. To generate *ptr-4(cs273)*, *ptr-4* single-guide RNA (AGGAATCGAAAGAACTGTGA) was injected with recombinant Cas9 protein (University of California, Berkeley) and marker pRF4 (*rol-6(su1006)*, after (Dokshin et al., 2018). Heterozygous *cs273* animals were recognized based on progeny embryonic lethality; the mutant allele was recovered over *szT1* and then rescued by a *ptr-4* transgene. *ptr-4(cs279)* arose during an attempted tagging experiment using the single-guide RNA that generated *ptr-4(cs264[PTR-4::SfGFP])* and the recombinant Cas9 protein and marker pRF4 used to generate *ptr-4(cs273)*; homozygous *cs279* animals were recognized based on their mild short and fat (Dpy) body shape. Edits in all alleles were determined by Sanger sequencing of genomic PCR products.

### RNAi

For Figs. 1 and 5, *ptr-4* double-stranded RNA (dsRNA) was synthesized using the Ambion MEGAscript RNAi Kit (AM1626; Ambion) and injected into gravid hermaphrodites at 6-12 ng/ul. Templates used were *ptr-4* RNAi bacterial clone C45B2.7 (Ahringer) (*ptr-4* RNAi 1) and genomic DNA amplified with primers oJC242 (TAATACGACTCACTATAGGGGTTTTCTTCAAATATGGAATATTTGTTG) and oJC243 (TAATACGACTCACTATAGGGCGTGGTTTTAATGATTTCCAGC) (*ptr-4* RNAi 2). For vulva aECM experiments, embryos laid 24–48 hr after injection were collected and observed 48-72 hrs later. Mock controls used dsRNA derived from the empty vector (EV) clone L4440 (Kamath and Ahringer, 2003). All other RNAi experiments were done by feeding animals bacteria expressing dsRNA (Timmons et al., 2001). *ptr-4*, *rab-5* and *rab-7* RNAi were carried out using sequenced and confirmed clones from the Ahringer library; as mock nontargeting dsRNA from the Ahringer library clone Y95B8A_84.g was used.

### Staging and microscopy

Embryos were selected at the 1.5-fold stage and then incubated for the number of hours indicated before mounting for imaging. L4 larvae were staged by vulva morphology. Live worms were immobilized with 50 mM levamisole in M9 and mounted on a slide with 2% agarose.

For experiments in Figs. 1, 3C,D, and 5, fluorescent and differential interference contrast (DIC) images were captured on a compound Zeiss Axioskop fitted with a Leica DFC360 FX camera, or with a Leica TCS SP8 confocal microscope. Images were processed and merged using ImageJ. To measure LET-653(ZP) fluorescence intensity, a 10-pixel-thick line was drawn across the vulE vulva region in L4.4 or L4.5 stage animals. The average fluorescence intensity of LET-653(ZP) in the three microns closest to the apical membrane peak fluorescence was divided by the average fluorescence of LET-653(ZP) in the center of the vulva to construct a ratio.

For experiments in Figs. 2 and 3A,B,E, worms were imaged with ORCA-flash 4.0 camera (Hamamatsu) mounted on an Axio Imager.M2 fluorescence microscope with a 63x Plan-Apochromat objective (Carl Zeiss, Germany) and a HXP 120 C light source using ZEN 2.6 software. All images were adjusted to the same parameters with OMERO.web 5.3.4-ice36-b69. For COL-19::GFP and DAPI staining, worms were imaged with an Olympus Fluoview FV3000 system with PTM voltage= 500, with an objective 60x 1.3NA with silicone oil, DAPI staining was performed according to Moribe et al., with minor modifications. For staining, we used 10 μg/μl of DAPI in M9. Animals were analyzed by fluorescence microscopy for DAPI staining in nuclei.

Structure illumination microscopy was done with DeltaVision OMX Optical Microscope (version 4), Software: DeltaVision OMX softWoRx. Oil 1.518 Images were analyzed in OMERO.web 5.3.4-ice36-b69.

### Swimming test

Young adults were placed on M9 buffer for 5 sec. The body bends were counted for 30 sec. At least eight worms of each condition were analyzed per experiment, and the experiment was performed three times. The results are presented as the average of tail movements per 30 sec. The data were analyzed with a one-tail ANOVA followed by Dunnett’s multiple comparisons test in Prism 7.

### Transmission election microscopy

For transmission electron microscopy (TEM) *C. elegans* animals were transferred to a droplet of M9 medium on a 100 μm cavity of a 3 mm aluminum specimen carrier (Engineering office M. Wohlwend GmbH, Sennwald, Switzerland). 5 - 10 worms were added to the droplet and the excess M9 medium was sucked off with dental filter tips. A flat aluminum specimen carrier was dipped in 1-hexadecene and added on top. Immediately, the specimen carrier sandwich was transferred to the middle plate of an HPM 100 high-pressure freezer (Leica Microsystems, Vienna, Austria) and frozen immediately without using ethanol as synchronizing medium. Freeze-substitution was carried out in water-free acetone containing 1% OsO_4_ for 8 hr at −90°C, 7 hr at ™60°C, 5 hr at −30°C, 1 hr at 0°C, with transition gradients of 30°C/hr, followed by 30 min incubation at RT. Samples were rinsed twice with acetone water-free, block-stained with 1% uranyl acetate in acetone (stock solution: 20% in MeOH) for 1 hr at 4°C, rinsed twice with water-free acetone and embedded in Epon/Araldite (Merck, Darmstadt, Germany): 66% in acetone overnight, 100% for 1 hr at RT and polymerized at 60°C for 20 hr. Ultrathin sections (50 nm) were post-stained with Reynolds lead citrate and imaged in a Talos 120 transmission electron microscope at 120 kV acceleration voltage equipped with a bottom mounted Ceta camera using the Maps software (Thermo Fisher Scientific, Eindhoven, The Netherlands).

## Acknowledgements

We gratefully acknowledge members of our laboratories for helpful comments and suggestions. The SIM microscopy was performed in the Imaging Core Facility of the Biozentrum. Some strains were provided by the Caenorhabditis Genetics Center (U. Minnesota), which is funded by the NIH Office of Research Resources (P40 OD010440). This work was funded by National Institutes of Health grants R01GM125959, R35GM136315 to M.V.S., T32 AR007465 to J.D.C, and by University of Basel (no grant number) and the Swiss National Science Foundation (310030_197779) to A.S.

---

**Figure S1.**
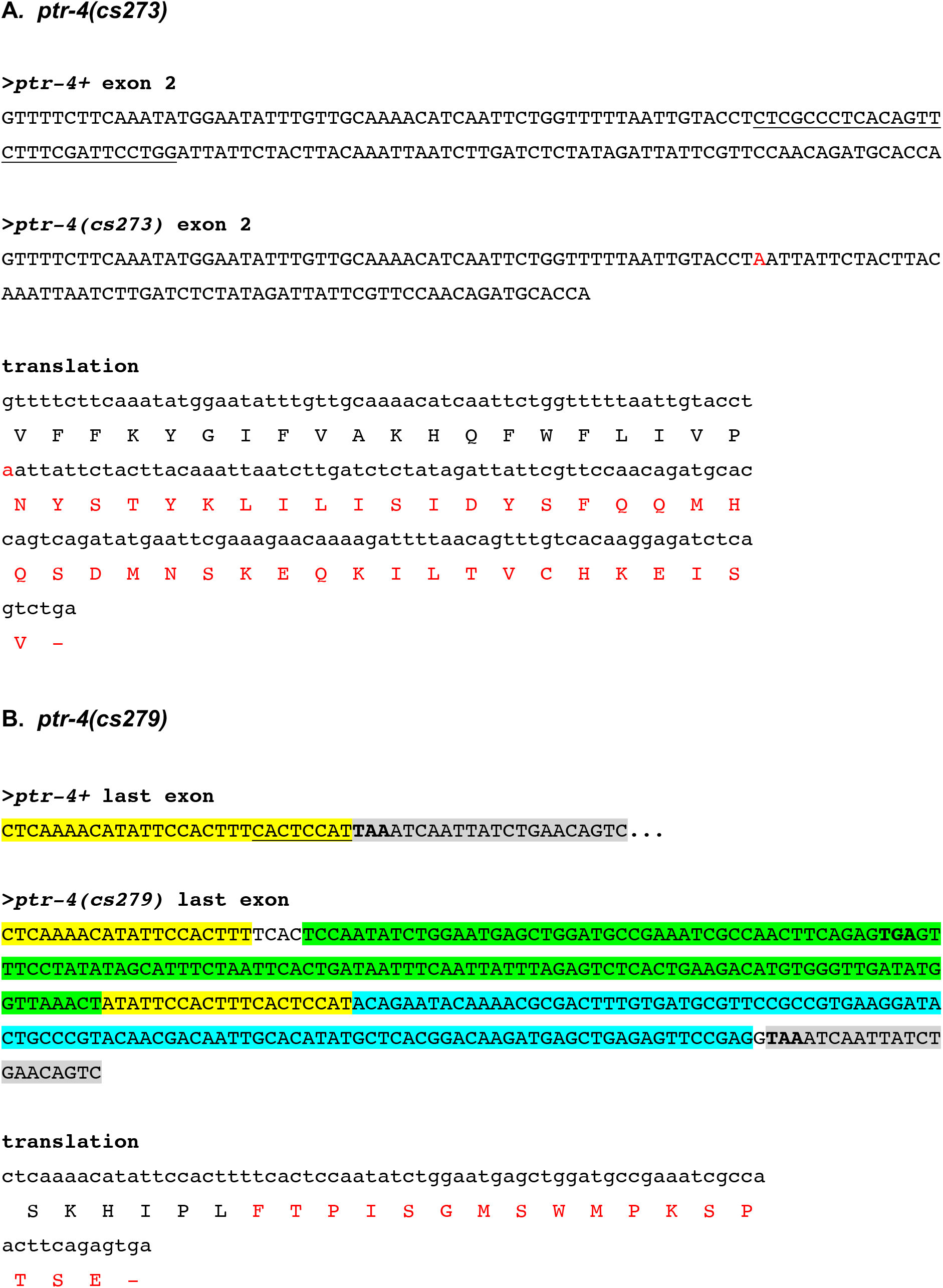
*ptr-4* allele sequences. (A) *ptr-4(cs273)* deletion. At top is the wild type *ptr-4* sequence from exon 2; the 29 nucleotides deleted in *cs273* are underlined. Below is the mutant sequence; an A (red) is inserted at the deletion site. Translation of this sequence yields a frameshift. (B) *ptr-4(cs279)* insertion/deletion. At top are 50 nucleotides of the wild type *ptr-4* sequence from the last exon; sequences disrupted by the insertion are underlined. Below is the mutant sequence. Yellow, *ptr-4* coding sequence. Grey, *ptr-4* stop codon and 3′ UTR. Green, *rol-6*-derived sequence. Aqua, *pie-1*-derived sequence. The newly introduced stop codon (TGA) and the endogenous stop codon (TAA) are indicated in bold font. Translation of this sequence yields a protein lacking the normal final 3 amino acids (SLH) and gaining seventeen aberrant residues (red). The majority of the inserted DNA comprises an extended 3′ UTR.

**Table S1:**
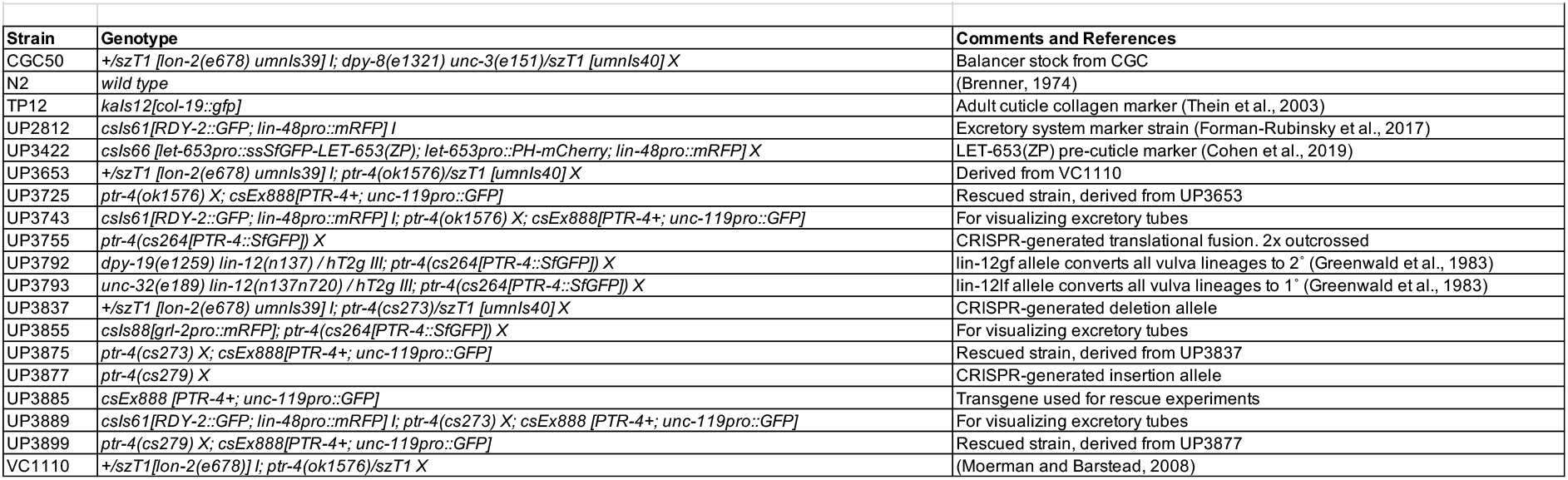
Strains used in this work.

